# A whole-brain model of auditory discrimination

**DOI:** 10.1101/2023.09.23.559095

**Authors:** A. Turan, E. Baspinar, A. Destexhe

## Abstract

Whole-brain simulations have been proposed previously to simulate global properties such as brain states or functional connectivity. Here, our aim is to build a whole-brain model to simulate a simple cognitive paradigm involving multiple brain areas. We focus on auditory discrimination, using a paradigm designed for the macaque cortex. To model at the whole-brain scale, we use The Virtual Brain (TVB) [18] simulation environment. TVB is a computational framework which simulates the brain as a network of small brain regions, where each node models neuronal populations and the connectivity between nodes determines the pathway of information flow over the brain. We use Adaptive Exponential (AdEx) neuronal population models [4, 11] to describe each node. For the connectivity, we use the open-access CoCoMac connectivity dataset [2], which is a matrix containing the connection weights between the nodes. We focus on a cognitive task that mainly involves the prefrontal cortex (PFC). In the auditory discrimination task, our pipeline starts from the primary auditory cortex stimulated by the auditory signals, it is then modulated in the PFC so that the stimulus discrimination occurs, after competition. Finally, it ends in the primary motor cortex which outputs the neuronal activity determining the motor action. Because the AdEx mean-fields can provide access to neuronal activity or local field potentials, we think that the present model constitutes a useful tool to promote interactions between theory and experiments for simple cognitive tasks in macaque monkey.

## 1 Introduction

Cognitive tasks at the whole-brain scale have been studied experimentally, with a lot of unknowns requiring strong computational support to be better understood. To contribute to this requirement, we propose a large-scale brain model which simulates the dynamics related to two auditory discrimination tasks. These tasks are based on distinguishing two auditory inputs having possibly different characteristics. To produce a cognitive behavior, the brain encodes and interprets sensory stimuli, weights evidence to select between alternatives, and finally, generates oriented motor actions. Our model considers this sensor-motor pathway at the whole-brain level in the macaque. This study follows from the model presented in [4, 5] and it is based on adapting this model to the TVB framework given in [13, 14].

Reliable measurement of neuronal activity is generally at the population level in neurophysiological experiments. It is obtained as averaged time-integrated population activity produced by the individual dynamics of the neurons in interaction within the population. The neuronal population can be modeled as a stochastic network system, which consists of a number of stochastic differential equations. Averaged network behavior models the population behavior. This requires high computational power since the network is high dimensional [23]. Another approach is to consider the averaged behavior of the network at the coarse-grained continuum limit where the number of neurons in the network is assumed to be infinity. In this way, the asymptotic limit of the network can be written in terms of the probability distribution of the state variables appearing in a single neuron in the network. This asymptotic limit is the so-called *mean-field limit* [1, 12, 22].

Adaptive Exponential (AdEx) mean-field framework [11, 24] approximates the neuronal population behavior modeled by the AdEx network [6]. In the case of the cerebral cortex, AdEx networks are used to model two cell types: Regular Spiking (RS) neurons, displaying spike-frequency adaptation as observed in the pyramidal (excitatory) neurons, and Fast Spiking (FS) neurons, with no adaptation, as observed in the interneurons (inhibitory). Hence, the AdEx networks are biophysically plausible. AdEx mean-field models are low dimensional, simpler and easier to analyze compared to the AdEx networks, yet they approximate closely the network dynamics, motivating our choice of model.

We proposed in [4, 5] a biophysically plausible AdEx mean-field model for decision-making related to a visual discrimination task applied on both human and macaque participants [9]. It was based on two AdEx mean-field models, each representing one single cortical column of PFC. In this setting, each column votes in favor of one of the two visual alternatives, where the winning vote is determined throughout a competition between the two columns. The competition is induced by intercolumnar excitation which is conflicted by intracolumnar inhibition. The excitation provokes the competition between the two columns. The winning column makes the decision. This competition is biased according to a function with two attractors. Each attractor promotes one of the two alternatives. This model is at mesoscale, more precisely, for only two columns of PFC. The stimulation was applied directly to PFC. The framework ignores the dynamics of the sensory (visual) cortex and motor areas, as well as the interactions of PFC with these regions. Therefore, it ignores the information routing in the brain, more specifically in the sensory-motor pathway corresponding to the relevant decision-making task.

To tackle these points, we use The Virtual Brain (TVB) [18], which is an open-source software and freely available on EBRAINS. It simulates all regions in the brain as a network of mean-field models. Each mean-field model approximates the dynamics of one region. We use the AdEx mean-field model embedded previously in TVB [13, 14]. This embedding was proposed to model awake and sleeping brain states, for both spontaneous and stimulated activity. However, it does not consider any cognitive task. Our principal contribution is to extend the AdEx mean-field model and its embedding in TVB such that TVB can produce the neuronal dynamics related to auditory discrimination. This can be considered as a first attempt towards modeling cognitive brain dynamics by using TVB.

The aim of this study is to simulate realistic brain dynamics of the two auditory discrimination tasks at the whole-brain scale. The motivation for choosing auditory tasks is that the relevant sensory-motor pathways are shorter in comparison to visual and somatosensory tasks. Challenges are both at connectivity and structural levels. At the connectivity level, introducing new cortical columns in the connectome requires introducing all connections between the new columns and all the other brain regions, as well as a careful normalization of the connectivity weights. At the structural level, information routing within the sensory-motor pathway requires an adequate encoding of the carried information between the regions in the pathway.

This work shows the potential of AdEx in simulating whole-brain dynamics corresponding to cognitive functions. Moreover, it provides computational support for experimental studies focusing on cognitive tasks in the macaque brain. It provides the whole-brain simulations based on AdEx, which is biophysically plausible, therefore can be tuned to the neural data recorded via multielectrode arrays.

In Section 2, we explain the auditory discrimination tasks considered in our simulations. In Section 3, we explain the TVB framework. Then, in Section 4, we describe the AdEx mean-field models which we use for the nodes in TVB. Finally, in Section 5, we present our TVB simulation results, and then, in Section 6, we provide the conclusion and perspectives.

## 2 Auditory discrimination tasks

We focus on two tasks. In the first task, the participant is provided sequentially two pitches with two possibly different frequencies. In the second task, these pitches are provided simultaneously. The participant is provided a button. The participant is instructed to push the button to indicate that it perceived the stimuli as different. Otherwise, the participant does not push the button, indicating that it perceived the stimuli as the same; see Figure 1 for an illustration of the tasks.

**Figure 1:**
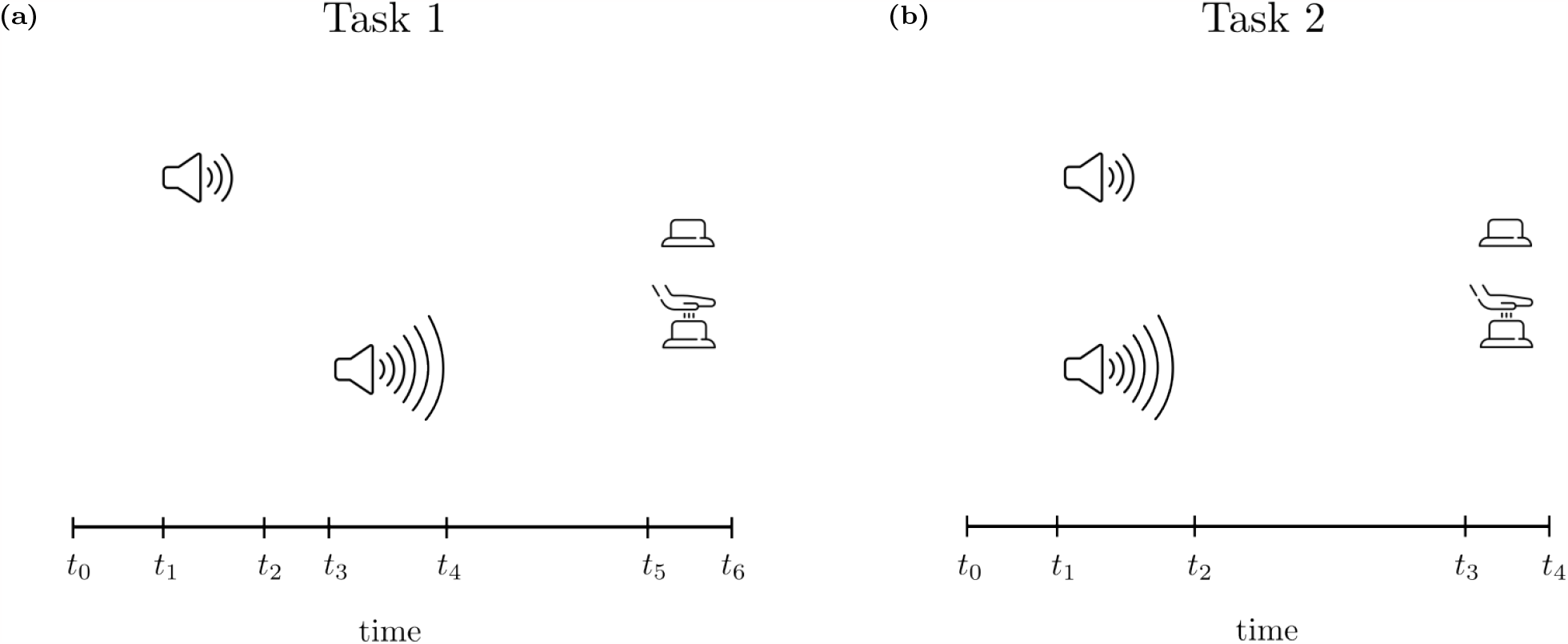
Auditory discrimination tasks. Task 1: The experiment starts at time *t*0. After an interval, the first stimulus is provided at time *t*_1_, and it lasts until *t*_2_. Then, there is no stimulus between *t*_2_ and *t*_3_. At *t*_3_, the second stimulus is provided, and it lasts until *t*_4_. After a waiting interval, starting from *t*_5_, the participant should push the button or not. Task 2: The same as Task 1. The only difference is that the stimuli are provided simultaneously between *t*_1_ and *t*_2_.

## 3 TVB

Large-scale brain simulators use parallel hardware to simulate a small brain region at the cellular level. This requires massive computational power and memory. TVB follows a different approach. It approximates the neuronal population dynamics via mean-field models of the population, thus decreases the complexity of micro-level mechanisms, allowing for macro-level simulations of the brain and/or the brain regions. It achieves such simplification at micro-level and such generalization at macro-level by merging the anatomical data from brain imaging data (diffusion MRI) with mean-field models of neuronal populations.

We have started from the TVB implementation based on the AdEx mean-field model proposed for the human brain [13, 14]. This implementation used the connectivity data regarding 129 regions of the human brain. This data was obtained during Human Connectome Project [20] by using diffusion MRI on the human brain. The connectome provided the connectivity weights of the long-range excitatory connections between all pairs which can be found among the 129 regions of the human brain. In this model, each region was represented as one single AdEx mean-field model [11]. This implementation was used to reproduce the neuronal dynamics of different brain states such as awake, sleep and anesthesia sates. In our study, we fix the parameters to remain always in the awake state.

We adapted the TVB-AdEx [13, 14] from the human brain to the monkey brain so as to study the neuronal dynamics induced by Tasks 1 and 2 at whole-brain level. In Task 1, we use the connectome of CoCoMac, which is for 84 regions of the macaque brain. Similarly to the human connectome, in the monkey connectome as well, we consider the long-range excitatory connections between regions. Each brain region is represented by one AdEx mean-field model [11]. In Task 2, we represent the dorsolateral PFC in terms of two AdEx mean-field models, which are connected to each other via long-range excitatory connections. This doubling is required to represent the competition between cortical columns in PFC, each corresponds to one of the two decisions. Moreover, the auditory cortex (A1) is organized in terms of columns, similarly to the other sensory cortices [10]. The tonotopic organization of A1 allows for pitch discrimination of the auditory stimuli. In [19], the segments of the input pattern are distributed via a gating module to separate the parts of PFC in an ordered manner. This results in a possible comparison of these segments with other segments coming from a second pattern [19], in coherence with the functional MRI experiments of an auditory recognition task. We model this gating by distributing the output of A1 selectively to different PFC areas (to PFCa and PFCb). This allows our model to initiate the discrimination task at A1 level in Task 2, as observed in the auditory discrimination experiments based on spatial selective detection of auditory stimuli [15].

The main difference of our TVB-AdEx implementation compared to the previous ones [13, 14] is that we introduce a simple cognitive context in the implementation by considering the whole sensory-motor pathway and by including the modulations of the neuronal dynamics as they propagate from one region to another.

## 4 AdEx model

We use the AdEx mean-field equations given in [11] to model the coarse-grained continuum activity of the neuronal population of each region. The AdEx framework models the averaged dynamics of excitatory (*e*) and inhibitory (*i*) populations, as well as their spike-frequency adaptation.

We follow the modeling hypothesis [4, 8, 9, 21], which is based on that a decision is the result of a competition between two pools (cortical columns), of neuronal populations. Each column represents one of the two alternatives. This hypothesis originates from two possible scenarios as mentioned in [8]. According to the first scenario, cells which are selective to a particular stimulus property are not spatially connected. Therefore, the connectivity would originate from Hebbian learning [7], in which the connection strength between two cells firing at the same time when a stimulus is presented would increase. On the contrary, long-term depression would weaken the connections between the cells which are not selective to the same stimulus property. In the second scenario, the cells which are selective to a particular stimulus property are not spatially distant from each other. They reside possibly in the same column. The connectivity is in coherence with the average distance between the cells. Two cells selective to the same stimulus property would be strongly connected, whereas two cells which are not selective to the same property would be distant from each other, therefore weakly or not connected. It is possible that the real cortex might be in an intermediate case, in which these scenarios participate together.

We use one single AdEx mean-field system for each region in the first auditory discrimination task. In this case, since the stimuli are provided sequentially, there is no need to use separate regions in PFC to model the dynamics encoding the two choices simultaneously. Consequently, there is no dynamic competition between any two regions of PFC, where each one of them votes in favor of one of the two choices: pushing the button or not pushing the button to indicate whether the stimuli were perceived as distinct or not, respectively. In the modeling of the second task, since the stimuli are provided simultaneously, we need two distinct regions in PFC, and possibly also in A1, to break the symmetry and provoke a dynamic competition resulting in the decision. To provoke the competition, we introduce an intercolumnar excitation which is conflicted by an intracolumnar inhibition.

For the second task, we used the model architecture provided in [4] for the two regions of PFC, see [4] for more details.

## 5 Simulations of the model

In our simulations, we modeled the difference of the pitches by changing the stimulus strength representing the auditory frequency. We introduced the stimuli to A1. We then analyzed the resulting data in different PFC areas of the macaque by using TVB. We aimed to discern which areas produced the most distinguishable signals. Regarding this point, the dorsolateral PFC (PFCdl) provided the most distinguishable results in our simulations. This is in coherence with the experimental evidence, which suggests the involvement of PFCdl in auditory discrimination tasks [3, 15–17]. Moreover, we took advantage of the hemispherical symmetry and restricted the simulations to the right hemisphere for the sake of simplicity.

Pipelines of the information flow in the simulations for Task 1 and Task 2 can be found in Figure 2. The simulation procedure for Task 1 can be summarized as follows: 1) Two stimuli are provided to A1 sequentially and independently. 2) The outputs of A1 are provided as inputs to PFCdl sequentially. 3) The difference between the outputs of PFCdl is measured based on mean squared error (MSE). 4) Depending on whether the MSE value is above or below a prefixed threshold, one of the two actions is chosen. Each action is modeled as a signal corresponding to a different signal strength. 5) The chosen action is transmitted to the motor cortex (M1) as the signal with the corresponding signal strength to the chosen action. The simulation procedure for Task 2 can be summarized as follows: 1) Two stimuli are provided to the duplicated A1 simultaneously.2) The outputs of the duplicated regions, A1-Ra and A1-Rb are checked if they exceed a prefixed threshold.If any of the output exceeds the threshold, then we provide a strong stimulus as input to the corresponding PFCdl region, otherwise a weak stimulus. 3)-5) are the same as in Task 1.

**Figure 2:**
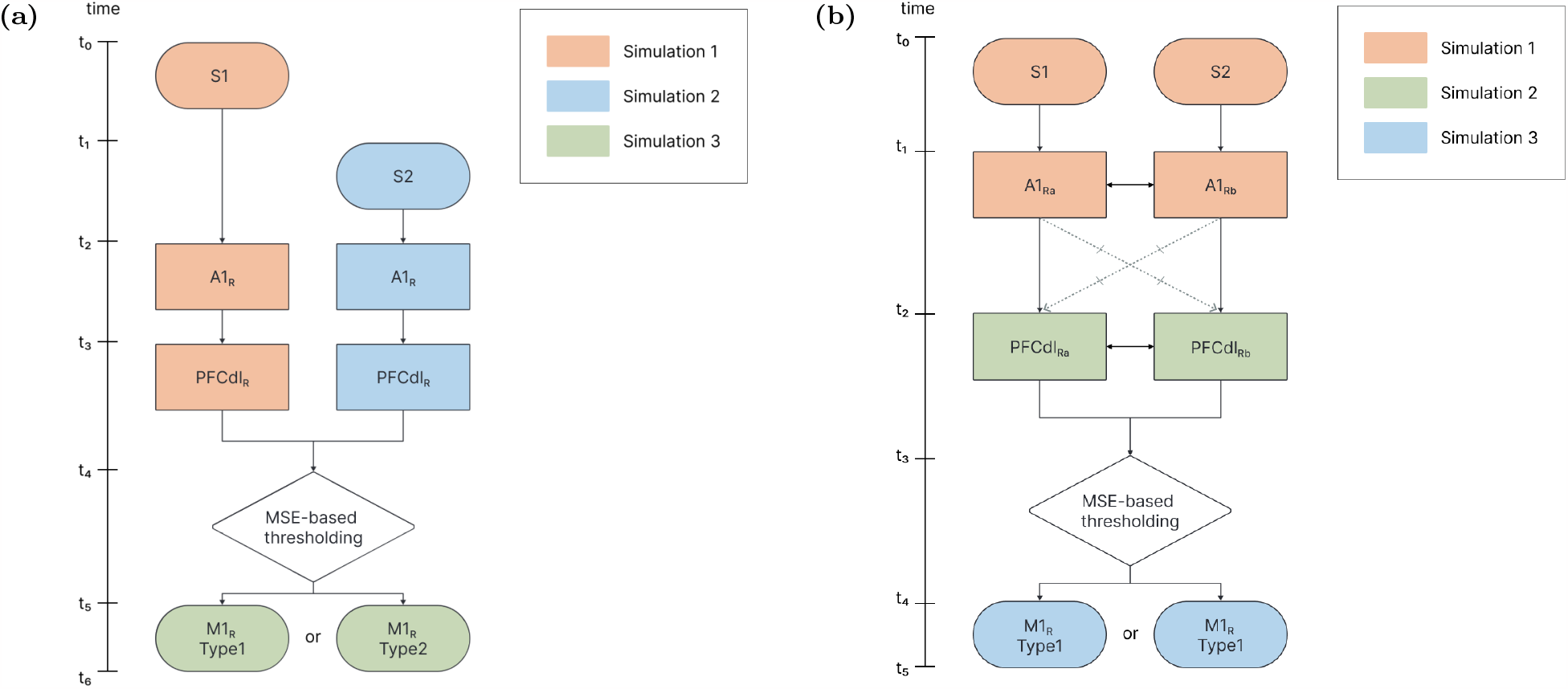
Information routing pipelines. (a) Task 1: We stimulate A1-Ra in the first simulation, and A1-Rb in the second. We compare the outputs of those simulations obtained from PFCdl-Ra and PFCdl-Rb via MSE. We perform the third simulation, which consists of stimulation of M1 with a stimulus generated according to the result of the comparison of the PFC outputs of the previous step. (b) Task 2: We stimulate A1 regions with two stimuli simultaneously in the first simulation. We check if the outputs of A1 regions exceed the prefixed threshold or not. In the second simulation, we stimulate PFC regions with strong and/or weak stimuli, depending on the thresholding result regarding the A1 outputs of the first simulation. The two PFC regions compete via their interconnections, and the result of this competition is compared using MSE. In the third simulation, we stimulate M1 with strong or weak stimulus, depending on the result of the MSE comparison.

### 5.1 Task 1

In this task, the stimuli are provided as (see Figure 1b-(a))

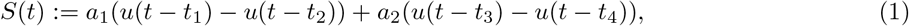

where *u* is the unit step function and *t*_2_*−t*_1_ = *t*_4_*−t*_3_. Here *a*_1_ and *a*_2_ represent the amplitudes of the first and second stimuli, respectively. One of the challenges was to determine the range of the stimulus strength which could yield to two distinguishable firing rate outputs in A1. Our findings reveal a pronounced change in the neuronal firing rate in PFCdl between the stimulus strength values of 0.405 and 0.410. In this way, we define two stimulus types: one with a stimulus strength below 0.405 (Type I), and the other above 0.410 (Type II).

Differences in neuronal responses to different stimuli allows for the categorization of the input signals in A1 into two groups: those associated with stimulus values less than or equal to 0.405 and those with stimulus values greater than or equal to 0.41. These findings allow for a thresholding to categorize the signals based on the stimulus strength. Therefore, by analyzing the output of PFCdl, we can determine retrospectively whether the auditory stimulus was Type I or Type II. For this purpose, we consider the inhibitory firing rate since the differences in the stimuli are manifested more clearly in inhibitory firing rates compared to the excitatory firing rates.

MSE value between the inhibitory reponses to any two stimuli stimX and stimY is denoted by

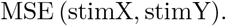

We name the stimuli given in Fig. 3 with values 0.001, 0.405, 0.410 and 2.000 as stim1, stim2, stim3 and stim4 separately. Some relevant MSE values are given as follows: MSE (stim1, stim2) = 34.57, MSE (stim3, stim4) = 2.73, MSE (stim2, stim3) = 91.22, MSE (stim1, stim4) = 123.35. A suitable threshold value for those MSE results is 60 for distinguishing between the inhibitory firing rates in PFCdl that were induced by different stimulus values. This threshold provides a reliable demarcation point, facilitating a clear differentiation of the neuronal responses based on their corresponding stimulus inputs. Depending on whether the MSE value exceeds 60 or not, we stimulate M1 by using an ad hoc stimulus, which is again a difference of two step functions scaled by a certain amplitude value similarly to the stimuli found in *S*(*t*) given by (1). The outputs from M1 which indicate that a motor action should be taken (push the button) or not are given in Figure 4.

**Figure 3:**
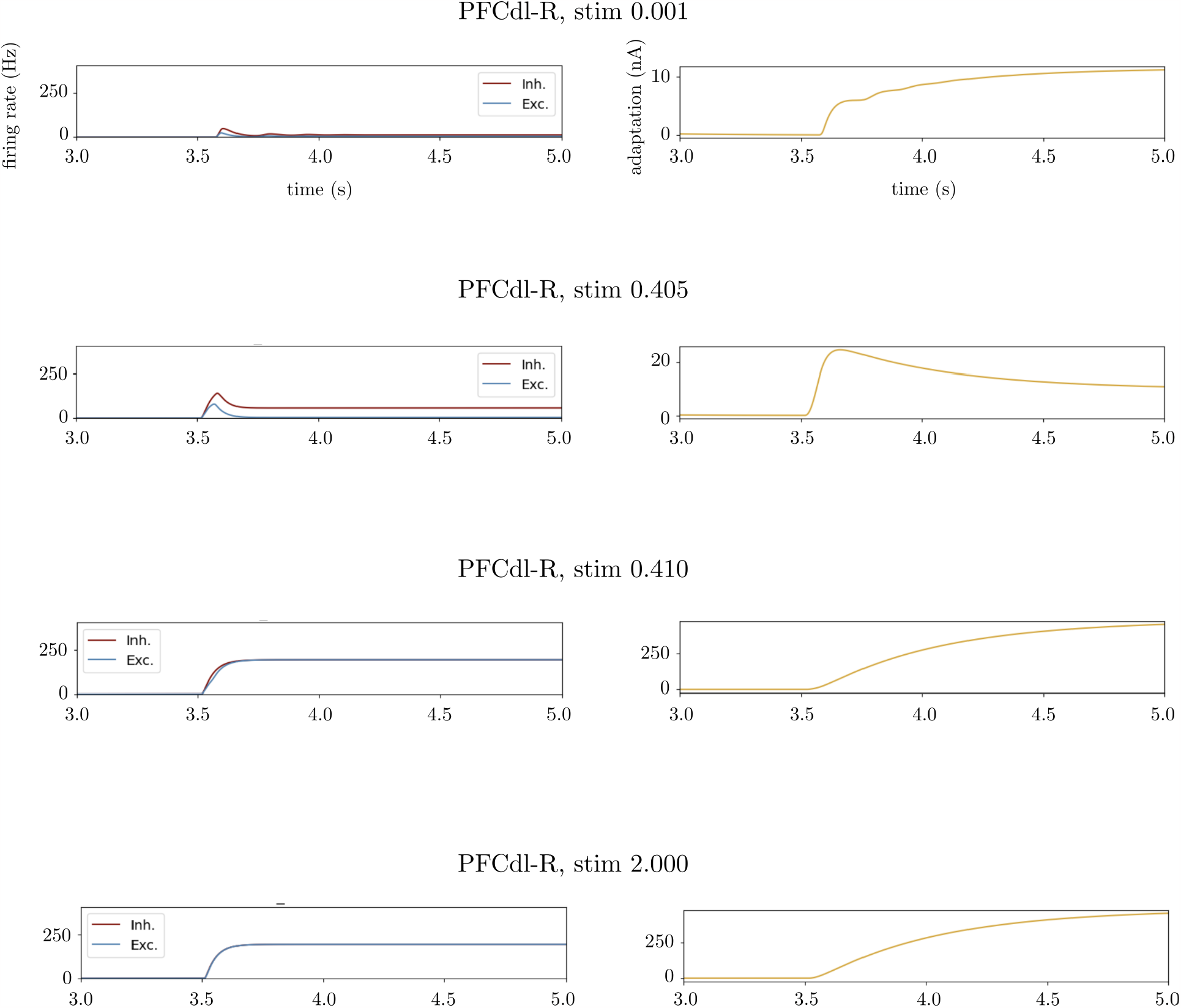
Firing rate and adaptation plots obtained from the dorsolateral PFC of the right hemisphere. The stimulus value varies from 0.001 to 2. The axes are shown in the first row and they are the same for all rows. The population behavior changes noticeably as the the stimulus value increases from 0.405 to 0.410.

**Figure 4:**
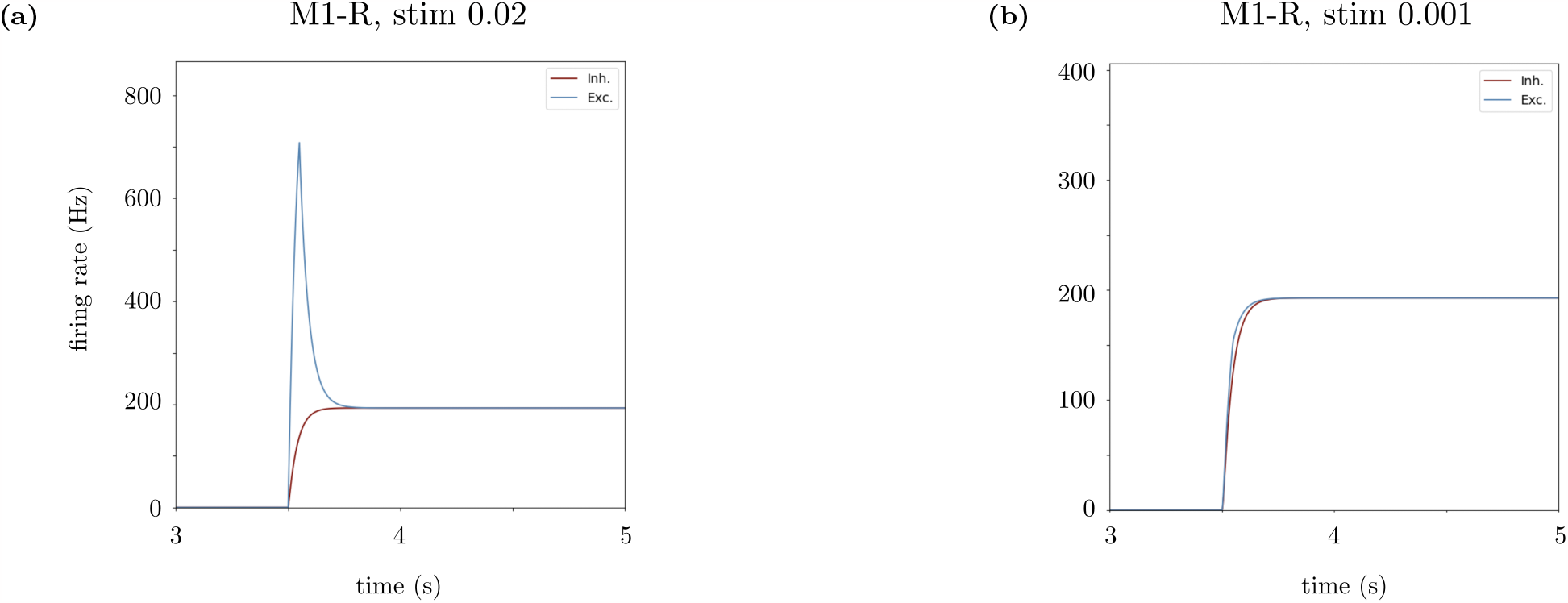
Outputs of the right hemisphere M1 area (M1-R) to two stimuli with amplitudes equal to 0.02 and 0.001. High amplitude stimulation in (a) results from a MSE value exceeding the threshold, thus it indicates that the model produces the activity for pushing the button. Small amplitude stimulation in (b) results from low MSE value, therefore M1-R activity is low, indicating no motor action.

### 5.2 Task 2

In the previous Task 1, two independent parallel streams were considered so there is no competition. In Task 2, we introduce a competition between the two streams via long-range excitatory connections between the corresponding cortical columns. In this task, the stimuli are provided as (see Figure 1b)

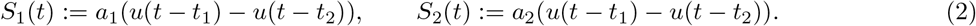

Another difference compared to Task 1 is that the stimuli are provided to the participant simultaneously, not sequentially. This requires two distinct regions in A1-R and PFCdl-R. In A1-R, each one of these regions is sensitive to one of the two stimuli. In PFCdl-R, each one of those two regions represents the neuronal populations voting in favor of one of the two decisions. After competition, if the stimuli are perceived as different then the decision is for pushing the button. Alternatively, if after competition, the stimuli are perceived as the same hence the decision is for not pushing the button.

To tackle this challenge, we introduced one additional region to each one of A1 and PFCdl separately. This results in two A1 and two PFCdl regions. We named these regions in A1 as A1a and A1b, and in PFCdl as PFCdla and PFCdlb. We obtained similar results to the ones of Task 1. We provide the outputs of the PFCdl regions induced by direct stimulations of PFCdla and PFCdlb in Figure 5 analogously to Figure 3. We present our simulation results corresponding to the pipeline given in Figure 2b in Figures 6 and 7.

**Figure 5:**
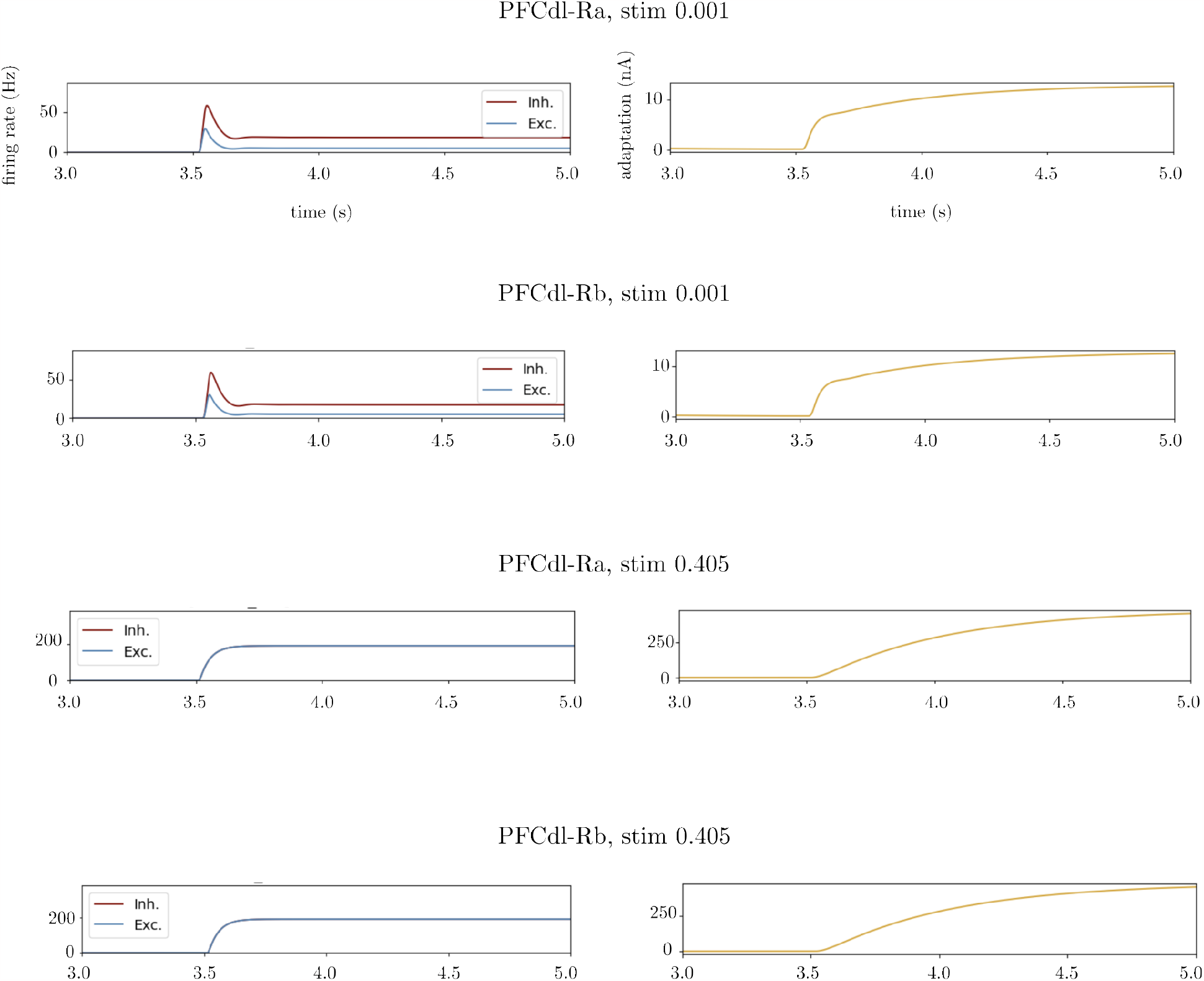
Firing rate and adaptation plots obtained from dorsolateral PFC of the right hemisphere, from both a and b regions. The stimulus value is changed from 0.001 to 0.405. The axes are shown in the first row and they are the same for all rows. Population behavior changes noticeably as the stimulus value increases from 0.001 to 0.405.

**Figure 6:**
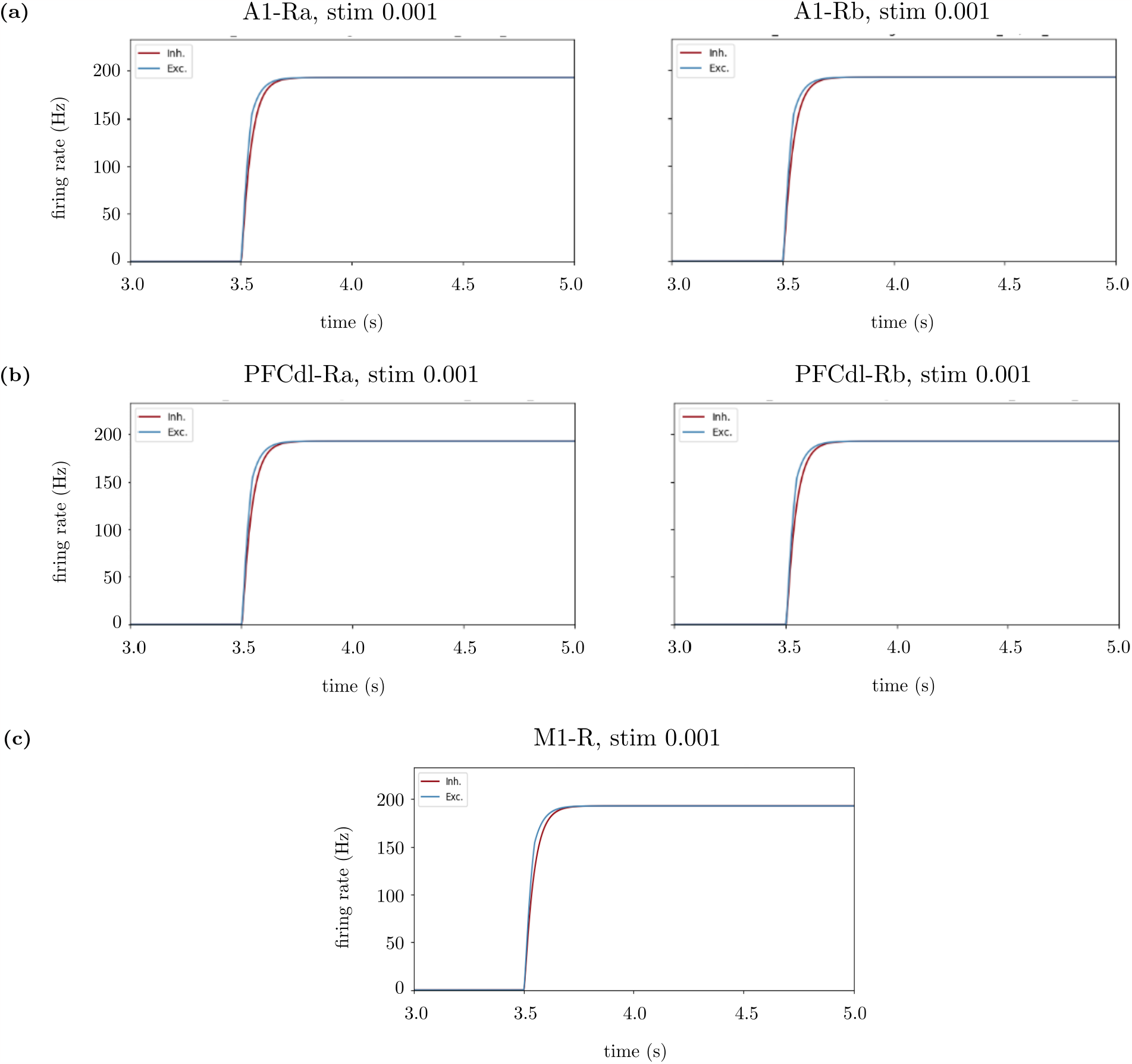
Simulation of Task 2 with two equivalent stimuli provided to A1. We provide the same pitches. The stimulus value is 0.001 for both. (a) Output firing rates from A1 regions. Depending on the output of each A1 region, we provide a strong or weak stimulus to the corresponding PFC region. (b) Output firing rates of the PFC regions. They are the same, meaning that the model perceive the stimuli as the same. Those outputs are compared to each other via the MSE thresholding; see Figure 2b. If MSE value is high, we provide a large stimulus to M1, meaning that the decision is for pushing the button, otherwise a weak stimulus is provided to M1. (c) Output of M1. It is small valued, so the decision is for not pushing the button.

**Figure 7:**
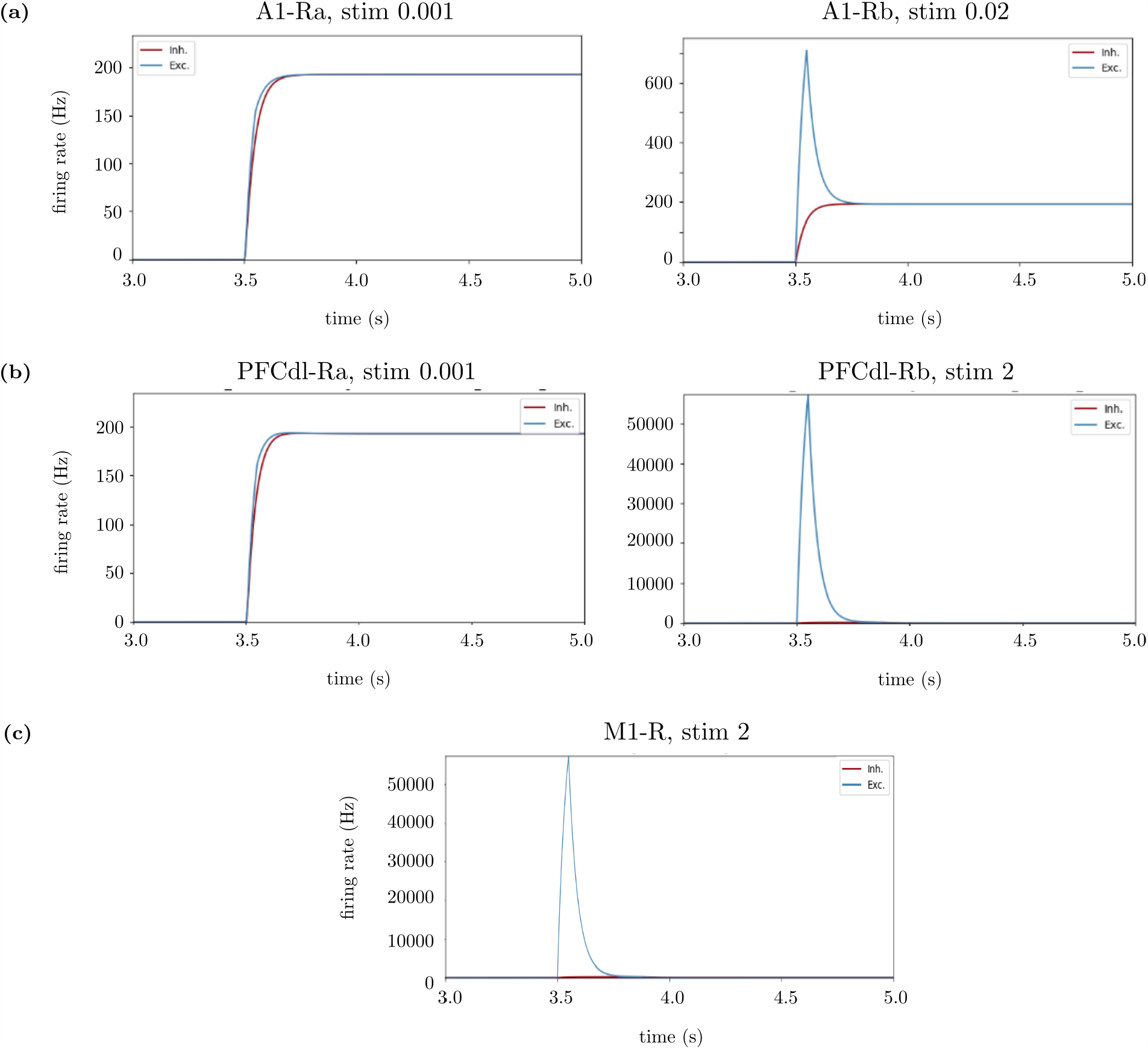
Simulation of Task 2 with two different stimuli provided to A1. We provide different pitches. The stimulus values are 0.001 and 0.02. (a) Output firing rates from A1 regions. Depending on the output of each A1 region, we provide a strong or weak stimulus to the corresponding PFC region. (b) Output firing rates of the PFC regions. PFCdl-Ra is stimulated by small stimulus and PFCdl-Rb is stimulated by large stimulus, meaning that the model perceive the stimuli as the different. Those outputs are compared to each other via the MSE thresholding; see Figure 2b. If MSE value is high, we provide a large stimulus to M1, meaning that the decision is for pushing the button, otherwise a weak stimulus is provided to M1. (c) Output of M1. It is large valued, so the decision is for pushing the button.

## 6 Conclusion

We provided a whole-brain scale model based on TVB integrated with a double-column AdEx mean-field framework. The model is for simulating neuronal dynamics of the brain regions relevant to simple auditory discrimination tasks, which can be considered as a first attempt of modeling of decision-making neuronal dynamics at the level of the whole brain.

The AdEx framework was already used for decision-making [4, 5]. However, this previous approach was at the level of neuronal populations constrained by the dynamics of the competition between two cortical columns. The input to the model was provided directly at PFC. Therefore, the information routing, or the interactions between principal brain regions involving in decision-making were not considered in this previous work. The novelty of the present work is that it extends this AdEx framework from the scale of the populations of a single brain region to multiple brain regions contributing to the relevant neuronal dynamics. We used a planning of the information routing, allowing to take into account the interactions between different brain regions.

It is important to note that in Task 2, there is a competition between the two input streams, and the decision (here the discrimination) is taken as a result of this competition, similar to the mechanism explored previously [4]. This results in a simulation that involves the routing of sensory information to the prefrontal cortex, where a decision is made, and the subsequent motor action evoked by the driving of M1 by PFC. The details of this information routing are probably over-simplified, but we believe they constitute a first step towards investigating cognitive paradigms by whole-brain models.

Another point of interest is the use of the AdEx mean-field framework, where the activities of excitatory and inhibitory populations, synaptic and membrane conductances, or local field potentials (LFPs), can be accessed. So in principle, this type of model can make predictions of the time course or respective timing of excitatory and inhibitory neurons in the different brain areas, the exact form and shape of LFPs, etc. These predictions can be tested by micro-electrode (or LFP) recordings in the corresponding areas in monkey. Thus, the present TVB-AdEx simulations provide a tool to promote bi-directional interactions between theory and experiments, and which can directly use electrophysiological recordings in monkey. We think that this tool should be very useful for a deeper understanding of the brain mechanisms underlying decision making, auditory discrimination and other cognitive paradigms.

## Supporting information

References

## 7 Acknowledgments

We hereby acknowledge that this research was supported by the Human Brain Project (European Union grant H2020-945539).

## References

[1] S.-i. Amari, Dynamics of pattern formation in lateral-inhibition type neural fields, Biological Cybernetics, 27 (1977), pp. 77–87.

[2] R. Bakker, T. Wachtler, and M. Diesmann, Cocomac 2.0 and the future of tract-tracing databases, Frontiers in neuroinformatics, 6 (2012), p. 30.

[3] H. Barbas, Architecture and cortical connections of the prefrontal cortex in the rhesus monkey, Advances in neurology, (1992), pp. 91–115.

[4] E. Baspinar, G. Cecchini, M. DePass, M. Andujar, P. Pani, S. Ferraina, R. Moreno-Bote,I. Cos, and A. Destexhe, A biologically plausible decision-making model based on interacting cortical columns, bioRxiv, (2023).

[5] E. Baspinar, G. Cecchini, R. Moreno-Bote, I. Cos, and A. Destexhe, Jupyter notebook of a biophysically plausible decision-making model based on interacting cortical columns, Zenodo, 2023.

[6] R. Brette and W. Gerstner, Adaptive exponential integrate-and-fire model as an effective description of neuronal activity, Journal of Neurophysiology, 94 (2005), pp. 3637–3642.

[7] N. Brunel, F. Carusi, and S. Fusi, Slow stochastic hebbian learning of classes of stimuli in a recurrent neural network, Network: Computation in Neural Systems, 9 (1998), p. 123.

[8] N. Brunel and X.-J. Wang, Effects of neuromodulation in a cortical network model of object working memory dominated by recurrent inhibition, Journal of Computational Neuroscience, 11 (2001), pp. 63–85.

[9] G. Cecchini, M. DePass, E. Baspinar, M. Andujar, S. Ramawat, P. Pani, S. Ferraina,A. Destexhe, R. Moreno-Bote, and I. Cos, A theoretical formalization of consequence-based decision-making, bioRxiv, (2023).

[10] D. F. Cechetto and J. C. Topolovec, Cerebral cortex, in Encyclopedia of the Human Brain, V. Ramachandran, ed., Academic Press, New York, 2002, pp. 663–679.

[11] M. Di Volo, A. Romagnoni, C. Capone, and A. Destexhe, Biologically realistic mean-field models of conductance-based networks of spiking neurons with adaptation, Neural Computation, 31 (2019), pp. 653–680.

[12] S. El Boustani and A. Destexhe, A master equation formalism for macroscopic modeling of asynchronous irregular activity states, Neural Computation, 21 (2009), pp. 46–100.

[13] J. S. Goldman, L. Kusch, D. Aquilue, B. H. YalçInkaya, D. Depannemaecker, K. Ancourt, T.-A. E. Nghiem, V. Jirsa, and A. Destexhe, A comprehensive neural simulation of slow-wave sleep and highly responsive wakefulness dynamics, Frontiers in Computational Neuroscience, 16 (2023).

[14] J. S. Goldman, L. Kusch, B. H. Yalcinkaya, D. Depannemaecker, T.-A. E. Nghiem, V. Jirsa, and A. Destexhe, Brain-scale emergence of slow-wave synchrony and highly responsive asynchronous states based on biologically realistic population models simulated in the virtual brain, bioRxiv, (2020).

[15] J. L. Napoli, C. R. Camalier, A.-L. Brown, J. Jacobs, M. M. Mishkin, and B. B. Averbeck, Correlates of auditory decision-making in prefrontal, auditory, and basal lateral amygdala cortical areas, Journal of Neuroscience, 41 (2021), pp. 1301–1316.

[16] M. Petrides and D. N. Pandya, Association fiber pathways to the frontal cortex from the superior temporal region in the rhesus monkey, Journal of Comparative Neurology, 273 (1988), pp. 52–66.

[17] B. Plakke and L. M. Romanski, Auditory connections and functions of prefrontal cortex, Frontiers in neuroscience, 8 (2014), p. 199.

[18] P. Sanz Leon, S. A. Knock, M. M. Woodman, L. Domide, J. Mersmann, A. R. McIntosh, and V. Jirsa, The virtual brain: a simulator of primate brain network dynamics, Frontiers in neuroinformatics, 7 (2013), p. 10.

[19] A. Ulloa, F. T. Husain, S. Kemeny, J. Xu, A. R. Braun, and B. Horwitz, Neural mechanisms of auditory discrimination of long-duration tonal patterns: a neural modeling and fmri study, Journal of integrative neuroscience, 7 (2008), pp. 501–527.

[20] D. C. Van Essen, K. Ugurbil, E. Auerbach, D. Barch, T. E. Behrens, R. Bucholz, A. Chang, L. Chen, M. Corbetta, S. W. Curtiss, et al., The human connectome project: a data acquisition perspective, Neuroimage, 62 (2012), pp. 2222–2231.

[21] X.-J. Wang, Probabilistic decision making by slow reverberation in cortical circuits, Neuron, 36 (2002), pp. 955–968.

[22] H. R. Wilson and J. D. Cowan, Excitatory and inhibitory interactions in localized populations of model neurons, Biophysical Journal, 12 (1972), pp. 1–24.

[23] Z.-C. Xiao, K. K. Lin, and L.-S. Young, A data-informed mean-field approach to mapping of cortical parameter landscapes, PLoS Computational Biology, 17 (2021), p. e1009718.

[24] Y. Zerlaut, S. Chemla, F. Chavane, and A. Destexhe, Modeling mesoscopic cortical dynamics using a mean-field model of conductance-based networks of adaptive exponential integrate-and-fire neurons, Journal of Computational Neuroscience, 44 (2018), pp. 45–61.

