## Supplementary material for "A whole-brain model of auditory discrimination": References: references.bib

\begin{thebibliography}{1}


@article{di2019biologically,
  title={Biologically realistic mean-field models of conductance-based networks of spiking neurons with adaptation},
  author={Di Volo, Matteo and Romagnoni, Alberto and Capone, Cristiano and Destexhe, Alain},
  journal={Neural Computation},
  volume={31},
  number={4},
  pages={653--680},
  year={2019},
  publisher={MIT Press}
}

@article{el2009master,
  title={A master equation formalism for macroscopic modeling of asynchronous irregular activity states},
  author={El Boustani, Sami and Destexhe, Alain},
  journal={Neural Computation},
  volume={21},
  number={1},
  pages={46--100},
  year={2009},
  publisher={MIT Press}
}

@article{zerlaut2018modeling,
  title={Modeling mesoscopic cortical dynamics using a mean-field model of conductance-based networks of adaptive exponential integrate-and-fire neurons},
  author={Zerlaut, Yann and Chemla, Sandrine and Chavane, Frederic and Destexhe, Alain},
  journal={Journal of Computational Neuroscience},
  volume={44},
  number={1},
  pages={45--61},
  year={2018},
  publisher={Springer}
}

@article{brette2005adaptive,
  title={Adaptive exponential integrate-and-fire model as an effective description of neuronal activity},
  author={Brette, Romain and Gerstner, Wulfram},
  journal={Journal of Neurophysiology},
  volume={94},
  number={5},
  pages={3637--3642},
  year={2005},
  publisher={American Physiological Society}
}

@article{finkelstein2021attractor,
  title={Attractor dynamics gate cortical information flow during decision-making},
  author={Finkelstein, Arseny and Fontolan, Lorenzo and Economo, Michael N and Li, Nuo and Romani, Sandro and Svoboda, Karel},
  journal={Nature Neuroscience},
  pages={1--8},
  year={2021},
  publisher={Nature Publishing Group}
}

@article{seidemann1998temporal,
  title={Temporal gating of neural signals during performance of a visual discrimination task},
  author={Seidemann, Eyal and Zohary, Ehud and Newsome, William T},
  journal={Nature},
  volume={394},
  number={6688},
  pages={72--75},
  year={1998},
  publisher={Nature Publishing Group}
}

@article{kiani2008bounded,
  title={Bounded integration in parietal cortex underlies decisions even when viewing duration is dictated by the environment},
  author={Kiani, Roozbeh and Hanks, Timothy D and Shadlen, Michael N},
  journal={Journal of Neuroscience},
  volume={28},
  number={12},
  pages={3017--3029},
  year={2008},
  publisher={Soc Neuroscience}
}

@article{kopec2015cortical,
  title={Cortical and subcortical contributions to short-term memory for orienting movements},
  author={Kopec, Charles D and Erlich, Jeffrey C and Brunton, Bingni W and Deisseroth, Karl and Brody, Carlos D},
  journal={Neuron},
  volume={88},
  number={2},
  pages={367--377},
  year={2015},
  publisher={Elsevier}
}

@article{van2018decoding,
  title={Decoding the influence of anticipatory states on visual perception in the presence of temporal distractors},
  author={Van Ede, Freek and Chekroud, Sammi R and Stokes, Mark G and Nobre, Anna C},
  journal={Nature Communications},
  volume={9},
  number={1},
  pages={1--12},
  year={2018},
  publisher={Nature Publishing Group}
}

@article{zuo2019rats,
  title={Rats generate vibrissal sensory evidence until boundary crossing triggers a decision},
  author={Zuo, Yanfang and Diamond, Mathew E},
  journal={Current Biology},
  volume={29},
  number={9},
  pages={1415--1424},
  year={2019},
  publisher={Elsevier}
}

@article{parker1998sense,
  title={Sense and the single neuron: probing the physiology of perception},
  author={Parker, Andrew J and Newsome, William T},
  journal={Annual Review of Neuroscience},
  volume={21},
  number={1},
  pages={227--277},
  year={1998},
  publisher={Annual Reviews 4139 El Camino Way, PO Box 10139, Palo Alto, CA 94303-0139, USA}
}

@article{romo2001touch,
  title={Touch and go: decision-making mechanisms in somatosensation},
  author={Romo, Ranulfo and Salinas, Emilio},
  journal={Annual Review of Neuroscience},
  volume={24},
  number={1},
  pages={107--137},
  year={2001},
  publisher={Annual Reviews 4139 El Camino Way, PO Box 10139, Palo Alto, CA 94303-0139, USA}
}

@article{schall2001neural,
  title={Neural basis of deciding, choosing and acting},
  author={Schall, Jeffrey D},
  journal={Nature Reviews Neuroscience},
  volume={2},
  number={1},
  pages={33--42},
  year={2001},
  publisher={Nature Publishing Group}
}

@article{broche2016neural,
  title={Neural substrates of decision-making},
  author={Broche-P{\'e}rez, Y and Jim{\'e}nez, LF Herrera and Omar-Mart{\'\i}nez, E},
  journal={Neurolog{\'\i}a (English Edition)},
  volume={31},
  number={5},
  pages={319--325},
  year={2016},
  publisher={Elsevier}
}

@article{paulus2003increased,
  title={Increased activation in the right insula during risk-taking decision making is related to harm avoidance and neuroticism},
  author={Paulus, Martin P and Rogalsky, Corianne and Simmons, Alan and Feinstein, Justin S and Stein, Murray B},
  journal={Neuroimage},
  volume={19},
  number={4},
  pages={1439--1448},
  year={2003},
  publisher={Elsevier}
}

@article{sanfey2006neuroeconomics,
  title={Neuroeconomics: cross-currents in research on decision-making},
  author={Sanfey, Alan G and Loewenstein, George and McClure, Samuel M and Cohen, Jonathan D},
  journal={Trends in Cognitive Sciences},
  volume={10},
  number={3},
  pages={108--116},
  year={2006},
  publisher={Elsevier}
}

@article{guo2013brain,
  title={Brain areas activated by uncertain reward-based decision-making in healthy volunteers},
  author={Guo, Zongjun and Chen, Juan and Liu, Shien and Li, Yuhuan and Sun, Bo and Gao, Zhenbo},
  journal={Neural Regeneration Research},
  volume={8},
  number={35},
  pages={3344},
  year={2013},
  publisher={Wolters Kluwer--Medknow Publications}
}

@article{albantakis2011changes,
  title={Changes of mind in an attractor network of decision-making},
  author={Albantakis, Larissa and Deco, Gustavo},
  journal={PLoS Comput Biol},
  volume={7},
  number={6},
  pages={e1002086},
  year={2011},
  publisher={Public Library of Science}
}

@article{moreno2007noise,
  title={Noise-induced alternations in an attractor network model of perceptual bistability},
  author={Moreno-Bote, Rub{\'e}n and Rinzel, John and Rubin, Nava},
  journal={Journal of Neurophysiology},
  volume={98},
  number={3},
  pages={1125--1139},
  year={2007},
  publisher={American Physiological Society}
}

@article{hayden2018neuronal,
  title={A neuronal theory of sequential economic choice},
  author={Hayden, Benjamin Y and Moreno-Bote, Rub{\'e}n},
  journal={Brain and Neuroscience Advances},
  volume={2},
  pages={2398212818766675},
  year={2018},
  publisher={SAGE Publications Sage UK: London, England}
}


@article{brunel2001effects,
  title={Effects of neuromodulation in a cortical network model of object working memory dominated by recurrent inhibition},
  author={Brunel, Nicolas and Wang, Xiao-Jing},
  journal={Journal of Computational Neuroscience},
  volume={11},
  number={1},
  pages={63--85},
  year={2001},
  publisher={Springer}
}

@article{wang2002probabilistic,
  title={Probabilistic decision making by slow reverberation in cortical circuits},
  author={Wang, Xiao-Jing},
  journal={Neuron},
  volume={36},
  number={5},
  pages={955--968},
  year={2002},
  publisher={Elsevier}
}


@article{marcos2013neural,
  title={Neural variability in premotor cortex is modulated by trial history and predicts behavioral performance},
  author={Marcos, Encarni and Pani, Pierpaolo and Brunamonti, Emiliano and Deco, Gustavo and Ferraina, Stefano and Verschure, Paul},
  journal={Neuron},
  volume={78},
  number={2},
  pages={249--255},
  year={2013},
  publisher={Elsevier}
}


@article{gilbert1979morphology,
  title={Morphology and intracortical projections of functionally characterised neurones in the cat visual cortex},
  author={Gilbert, Charles D and Wiesel, Torsten N},
  journal={Nature},
  volume={280},
  number={5718},
  pages={120--125},
  year={1979},
  publisher={Nature Publishing Group}
}

@article{rockland1982widespread,
  title={Widespread periodic intrinsic connections in the tree shrew visual cortex},
  author={Rockland, Kathleen S and Lund, Jennifer S},
  journal={Science},
  volume={215},
  number={4539},
  pages={1532--1534},
  year={1982},
  publisher={American Association for the Advancement of Science}
}

@article{livingstone1984specificity,
  title={Specificity of intrinsic connections in primate primary visual cortex},
  author={Livingstone, MARGARET S and Hubel, DAVID H},
  journal={Journal of Neuroscience},
  volume={4},
  number={11},
  pages={2830--2835},
  year={1984},
  publisher={Soc Neuroscience}
}

@article{martin1984form,
  title={Form, function and intracortical projections of spiny neurones in the striate visual cortex of the cat.},
  author={Martin, KA and Whitteridge, David},
  journal={The Journal of Physiology},
  volume={353},
  number={1},
  pages={463--504},
  year={1984},
  publisher={Wiley Online Library}
}

@article{kisvarday1986synaptic,
  title={Synaptic targets of HRP-filled layer III pyramidal cells in the cat striate cortex},
  author={Kisv{\'a}rday, Zolt{\'a}n F and Martin, Kevan AC and Freund, TF and Magloczky, ZS and Whitteridge, D and Somogyi, P},
  journal={Experimental Brain Research},
  volume={64},
  number={3},
  pages={541--552},
  year={1986},
  publisher={Springer}
}

@article{mcguire1991targets,
  title={Targets of horizontal connections in macaque primary visual cortex},
  author={McGuire, Barbara A and Gilbert, Charles D and Rivlin, Patricia K and Wiesel, Torsten N},
  journal={Journal of Comparative Neurology},
  volume={305},
  number={3},
  pages={370--392},
  year={1991},
  publisher={Wiley Online Library}
}

@article{albantakis2009encoding,
  title={The encoding of alternatives in multiple-choice decision making},
  author={Albantakis, Larissa and Deco, Gustavo},
  journal={Proceedings of the National Academy of Sciences},
  volume={106},
  number={25},
  pages={10308--10313},
  year={2009},
  publisher={National Acad Sciences}
}

@article{churchland2008decision,
  title={Decision-making with multiple alternatives},
  author={Churchland, Anne K and Kiani, Roozbeh and Shadlen, Michael N},
  journal={Nature Neuroscience},
  volume={11},
  number={6},
  pages={693--702},
  year={2008},
  publisher={Nature Publishing Group}
}

@article{arnold1974stochastic,
  title={Stochastic differential equations},
  author={Arnold, Ludwig},
  journal={New York},
  year={1974}
}

@article{goldman2020brain,
  title={Brain-scale emergence of slow-wave synchrony and highly responsive asynchronous states based on biologically realistic population models simulated in The Virtual Brain},
  author={Goldman, Jennifer S and Kusch, Lionel and Yalcinkaya, Bahar Hazal and Depannemaecker, Damien and Nghiem, Trang-Anh E and Jirsa, Viktor and Destexhe, Alain},
  journal={bioRxiv},
  year={2020},
  publisher={Cold Spring Harbor Laboratory}
}

@article{drugowitsch2012cost,
  title={The cost of accumulating evidence in perceptual decision making},
  author={Drugowitsch, Jan and Moreno-Bote, Rub{\'e}n and Churchland, Anne K and Shadlen, Michael N and Pouget, Alexandre},
  journal={Journal of Neuroscience},
  volume={32},
  number={11},
  pages={3612--3628},
  year={2012},
  publisher={Soc Neuroscience}
}

@article{mochol2021prefrontal,
  title={Prefrontal cortex represents heuristics that shape choice bias and its integration into future behavior},
  author={Mochol, Gabriela and Kiani, Roozbeh and Moreno-Bote, Rub{\'e}n},
  journal={Current Biology},
  volume={31},
  number={6},
  pages={1234--1244},
  year={2021},
  publisher={Elsevier}
}

@article{shadlen2013decision,
  title={Decision making as a window on cognition},
  author={Shadlen, Michael N and Kiani, Roozbeh},
  journal={Neuron},
  volume={80},
  number={3},
  pages={791--806},
  year={2013},
  publisher={Elsevier}
}

@article{rushworth2011frontal,
  title={Frontal cortex and reward-guided learning and decision-making},
  author={Rushworth, Matthew FS and Noonan, MaryAnn P and Boorman, Erie D and Walton, Mark E and Behrens, Timothy E},
  journal={Neuron},
  volume={70},
  number={6},
  pages={1054--1069},
  year={2011},
  publisher={Elsevier}
}

@article{domenech2015executive,
  title={Executive control and decision-making in the prefrontal cortex},
  author={Domenech, Philippe and Koechlin, Etienne},
  journal={Current Opinion in Behavioral Sciences},
  volume={1},
  pages={101--106},
  year={2015},
  publisher={Elsevier}
}

@article{bechara2000characterization,
  title={Characterization of the decision-making deficit of patients with ventromedial prefrontal cortex lesions},
  author={Bechara, Antoine and Tranel, Daniel and Damasio, Hanna},
  journal={Brain},
  volume={123},
  number={11},
  pages={2189--2202},
  year={2000},
  publisher={Oxford University Press}
}

@article{hampton2006role,
  title={The role of the ventromedial prefrontal cortex in abstract state-based inference during decision making in humans},
  author={Hampton, Alan N and Bossaerts, Peter and O’doherty, John P},
  journal={Journal of Neuroscience},
  volume={26},
  number={32},
  pages={8360--8367},
  year={2006},
  publisher={Soc Neuroscience}
}

@article{sirigu2022reward,
  title={Reward and decision processes in the brains of humans and nonhuman primates},
  author={Sirigu, Angela and Duhamel, Jean-Ren{\'e}},
  journal={Dialogues in Clinical Neuroscience},
  year={2022},
  publisher={Taylor \& Francis}
}

@article{monosov2012regionally,
  title={Regionally distinct processing of rewards and punishments by the primate ventromedial prefrontal cortex},
  author={Monosov, Ilya E and Hikosaka, Okihide},
  journal={Journal of Neuroscience},
  volume={32},
  number={30},
  pages={10318--10330},
  year={2012},
  publisher={Soc Neuroscience}
}

@article{suzuki2021distinct,
  title={Distinct regions of the striatum underlying effort, movement initiation and effort discounting},
  author={Suzuki, Shosuke and Lawlor, Victoria M and Cooper, Jessica A and Arulpragasam, Amanda R and Treadway, Michael T},
  journal={Nature human behaviour},
  volume={5},
  number={3},
  pages={378--388},
  year={2021},
  publisher={Nature Publishing Group UK London}
}

@article{xiao2021data,
  title={A data-informed mean-field approach to mapping of cortical parameter landscapes},
  author={Xiao, Zhuo-Cheng and Lin, Kevin K and Young, Lai-Sang},
  journal={PLoS Computational Biology},
  volume={17},
  number={12},
  pages={e1009718},
  year={2021},
  publisher={Public Library of Science San Francisco, CA USA}
}

@article{wilson1972excitatory,
  title={Excitatory and inhibitory interactions in localized populations of model neurons},
  author={Wilson, Hugh R and Cowan, Jack D},
  journal={Biophysical Journal},
  volume={12},
  number={1},
  pages={1--24},
  year={1972},
  publisher={Elsevier}
}

@article{amari1977dynamics,
  title={Dynamics of pattern formation in lateral-inhibition type neural fields},
  author={Amari, Shun-ichi},
  journal={Biological Cybernetics},
  volume={27},
  number={2},
  pages={77--87},
  year={1977},
  publisher={Springer}
}


@book{doya2007bayesian,
  title={Bayesian brain: Probabilistic approaches to neural coding},
  author={Doya, Kenji and Ishii, Shin and Pouget, Alexandre and Rao, Rajesh PN},
  year={2007},
  publisher={MIT press}
}

@article{giamundo2021neuronal,
  title={Neuronal activity in the premotor cortex of monkeys reflects both cue salience and motivation for action generation and inhibition},
  author={Giamundo, Margherita and Giarrocco, Franco and Brunamonti, Emiliano and Fabbrini, Francesco and Pani, Pierpaolo and Ferraina, Stefano},
  journal={Journal of Neuroscience},
  volume={41},
  number={36},
  pages={7591--7606},
  year={2021},
  publisher={Soc Neuroscience}
}

@article{leotti2010motivational,
  title={Motivational influences on response inhibition measures.},
  author={Leotti, Lauren A and Wager, Tor D},
  journal={Journal of Experimental Psychology: Human Perception and Performance},
  volume={36},
  number={2},
  pages={430},
  year={2010},
  publisher={American Psychological Association}
}

@article{padmala2010interactions,
  title={Interactions between cognition and motivation during response inhibition},
  author={Padmala, Srikanth and Pessoa, Luiz},
  journal={Neuropsychologia},
  volume={48},
  number={2},
  pages={558--565},
  year={2010},
  publisher={Elsevier}
}

@software{baspinarJupyter2023,
  author       = {Emre Baspinar and
                  Gloria Cecchini and
                  Ruben Moreno-Bote and
                  Ignasi Cos and
                  Alain Destexhe},
  title        = {{Jupyter notebook of a biophysically plausible 
                   decision-making model based on interacting
                   cortical columns}},
  month        = {\textit{Zenodo},},
  year         = 2023,
  publisher    = {Zenodo},
  version      = {1.0.0},
  doi          = {10.5281/zenodo.7682309},
  url          = {https://doi.org/10.5281/zenodo.7682309},
  journal       = {Zenodo}
}

@article {cecchiniPreprint,
	author = {Cecchini, Gloria and DePass, Michael and Baspinar, Emre and Andujar, Marta and Ramawat, Surabhi and Pani, Pierpaolo and Ferraina, Stefano and Destexhe, Alain and Moreno-Bote, Rub{\'e}n and Cos, Ignasi},
	title = {A Theoretical Formalization of Consequence-Based Decision-Making},
	elocation-id = {2023.02.14.528595},
	year = {2023},
	doi = {10.1101/2023.02.14.528595},
	publisher = {Cold Spring Harbor Laboratory},
	URL = {https://www.biorxiv.org/content/early/2023/02/15/2023.02.14.528595},
	eprint = {https://www.biorxiv.org/content/early/2023/02/15/2023.02.14.528595.full.pdf},
	journal = {bioRxiv}
}

@article{sanz2013virtual,
  title={The Virtual Brain: a simulator of primate brain network dynamics},
  author={Sanz Leon, Paula and Knock, Stuart A and Woodman, M Marmaduke and Domide, Lia and Mersmann, Jochen and McIntosh, Anthony R and Jirsa, Viktor},
  journal={Frontiers in neuroinformatics},
  volume={7},
  pages={10},
  year={2013},
  publisher={Frontiers Media SA}
}


@ARTICLE{goldman2021comprehensive,  
AUTHOR={Goldman, Jennifer S. and Kusch, Lionel and Aquilue, David and Yalçınkaya, Bahar Hazal and Depannemaecker, Damien and Ancourt, Kevin and Nghiem, Trang-Anh E. and Jirsa, Viktor and Destexhe, Alain},   
	 
TITLE={A comprehensive neural simulation of slow-wave sleep and highly responsive wakefulness dynamics},      
	
JOURNAL={Frontiers in Computational Neuroscience},      
	
VOLUME={16},           
	
YEAR={2023},      
	  
URL={https://www.frontiersin.org/articles/10.3389/fncom.2022.1058957},       
	
DOI={10.3389/fncom.2022.1058957},      
	
ISSN={1662-5188},     

}


@article {baspinar2023biologically,
	author = {E. Baspinar and G. Cecchini and M. DePass and M. Andujar and P. Pani and S. Ferraina and R. Moreno-Bote and I. Cos and A. Destexhe},
	title = {A biologically plausible decision-making model based on interacting cortical columns},
	elocation-id = {2023.02.28.530384},
	year = {2023},
	doi = {10.1101/2023.02.28.530384},
	publisher = {Cold Spring Harbor Laboratory},
	URL = {https://www.biorxiv.org/content/early/2023/03/01/2023.02.28.530384},
	eprint = {https://www.biorxiv.org/content/early/2023/03/01/2023.02.28.530384.full.pdf},
	journal = {bioRxiv}
}

@article{ulloa2008neural,
  title={Neural mechanisms of auditory discrimination of long-duration tonal patterns: a neural modeling and fMRI study},
  author={Ulloa, Antonio and Husain, Fatima T and Kemeny, Stefan and Xu, Jiang and Braun, Allen R and Horwitz, Barry},
  journal={Journal of integrative neuroscience},
  volume={7},
  number={04},
  pages={501--527},
  year={2008},
  publisher={World Scientific}
}

@incollection{CECHETTO2002663,
title = {Cerebral Cortex},
editor = {V.S. Ramachandran},
booktitle = {Encyclopedia of the Human Brain},
publisher = {Academic Press},
address = {New York},
pages = {663-679},
year = {2002},
isbn = {978-0-12-227210-3},
doi = {https://doi.org/10.1016/B0-12-227210-2/00087-X},
url = {https://www.sciencedirect.com/science/article/pii/B012227210200087X},
author = {David F. Cechetto and Jane C. Topolovec}
}


@article{napoli2021correlates,
  title={Correlates of auditory decision-making in prefrontal, auditory, and basal lateral amygdala cortical areas},
  author={Napoli, Julia L and Camalier, Corrie R and Brown, Anna-Leigh and Jacobs, Jessica and Mishkin, Mortimer M and Averbeck, Bruno B},
  journal={Journal of Neuroscience},
  volume={41},
  number={6},
  pages={1301--1316},
  year={2021},
  publisher={Soc Neuroscience}
}

@article{brunel1998slow,
  title={Slow stochastic Hebbian learning of classes of stimuli in a recurrent neural network},
  author={Brunel, Nicolas and Carusi, Francesco and Fusi, Stefano},
  journal={Network: Computation in Neural Systems},
  volume={9},
  number={1},
  pages={123},
  year={1998},
  publisher={IOP Publishing}
}

@article{bakker2012cocomac,
  title={CoCoMac 2.0 and the future of tract-tracing databases},
  author={Bakker, Rembrandt and Wachtler, Thomas and Diesmann, Markus},
  journal={Frontiers in neuroinformatics},
  volume={6},
  pages={30},
  year={2012},
  publisher={Frontiers Media SA}
}

@article{van2012human,
  title={The Human Connectome Project: a data acquisition perspective},
  author={Van Essen, David C and Ugurbil, Kamil and Auerbach, Edward and Barch, Deanna and Behrens, Timothy EJ and Bucholz, Richard and Chang, Acer and Chen, Liyong and Corbetta, Maurizio and Curtiss, Sandra W and others},
  journal={Neuroimage},
  volume={62},
  number={4},
  pages={2222--2231},
  year={2012},
  publisher={Elsevier}
}

@article{plakke2014auditory,
  title={Auditory connections and functions of prefrontal cortex},
  author={Plakke, Bethany and Romanski, Lizabeth M},
  journal={Frontiers in neuroscience},
  volume={8},
  pages={199},
  year={2014},
  publisher={Frontiers Media SA}
}

@article{petrides1988association,
  title={Association fiber pathways to the frontal cortex from the superior temporal region in the rhesus monkey},
  author={Petrides, Michael and Pandya, Deepak N},
  journal={Journal of Comparative Neurology},
  volume={273},
  number={1},
  pages={52--66},
  year={1988},
  publisher={Wiley Online Library}
}

@article{barbas1992architecture,
  title={Architecture and cortical connections of the prefrontal cortex in the rhesus monkey},
  author={Barbas, H},
  journal={Advances in neurology},
  pages={91--115},
  year={1992},
  publisher={Raven Press}
}


\end{thebibliography}
